# Tertiary lymphoid structure as a therapeutic target of Crohn’s disease: modulation by TNFα blockade

**DOI:** 10.1101/2024.11.23.625032

**Authors:** Yoshihiro Nagase, Soh Okano, Asuka Furukawa, Takumi Kanazawa, Kouhei Yamamoto, Kurara Yamamoto, Kenichi Ohashi, Tetsuo Yamana, Shuko Nojiri, Yutaka Suzuki, Akinori Kanai, Takashi Yao, Keiko Abe, Toshiaki Ohteki, Makoto Kodama

**Author notes:** Corresponding author: Makoto Kodama Department of Pathology, Tokyo Yamate Medical Center 3-22-1 Hyakunincho, Shinjuku-Ku, Tokyo, Japan. Contributed equally (Yoshihiro Nagase, Makoto Kodama).

## Abstract

Inflammatory diseases, such as Crohn’s disease (CD), are primarily treated with therapies aimed at alleviating inflammation. Notably, biological agents like TNFα inhibitors have transformed the management of these conditions. Previous research has indicated that TNFα inhibitors affect the cells in the lamina propria. However, this study demonstrates that infliximab, a type of TNFα inhibitor, reduces the presence of tertiary lymphoid structures (TLS) and alleviates inflammation of CD by targeting follicular dendritic cells and macrophages within these structures. TLS could also be a potential target for various biological agents used to treat inflammatory diseases beyond CD. Furthermore, our study suggests that the status of TLS may serve as a predictor for the response to TNFα inhibitor treatment. Our research could pave the way for new treatment strategies and the advancement of personalized medicine for inflammatory diseases.

## INTRODUCTION

Crohn’s disease (CD) is an inflammatory bowel disease of unknown etiology that causes chronic inflammation and ulceration of the intestinal tract. As the cause of the disease remains unclear, treatment focuses on alleviating symptoms and suppressing inflammation. Notably, the introduction of biological agents for inflammatory bowel diseases, specifically TNFα inhibitors that target inflammatory cytokines, has led to high rates of remission induction and has significantly transformed the treatment approach for CD^1^. Regarding the mechanism of action of TNFα inhibitors such as Infliximab (IFX), numerous studies have demonstrated how the inhibition of TNFα contributes to a reduction in inflammation. Specifically, IFX alleviates inflammation through several processes. It neutralizes both soluble TNF (sTNF) and transmembrane TNF (mTNF), induces apoptosis in mTNF- expressing cells, promotes T-cell apoptosis via mTNF-expressing CD14 macrophages, and encourages macrophages to adopt a wound-healing phenotype (M2 phenotype) by interacting with the Fc receptor^2,3^. Several reports have indicated that TNFα inhibitors affect T cells and macrophages in the lamina propria^2,3^. However, IFX-induced apoptosis of cells in the lamina propria may be a secondary effect related to the resolution of the primary inflammatory process^4^. In terms of the prominent inflammation, a well-known pattern in CD is known as disproportionate inflammation, which is more severe in the submucosal layer or more profound than in the lamina propria^5^. This means that the significant inflammation seen in CD is not primarily located in the lamina propria, where the cells targeted by IFX are thought to be found, but instead occurs in the submucosal layer or deeper (Fig. 1a). To fully understand the mechanism of action of IFX, it is essential to evaluate changes throughout the intestinal wall, particularly in areas with the highest levels of inflammation, following IFX treatment using surgical specimens from patients with CD.

**Fig. 1.**
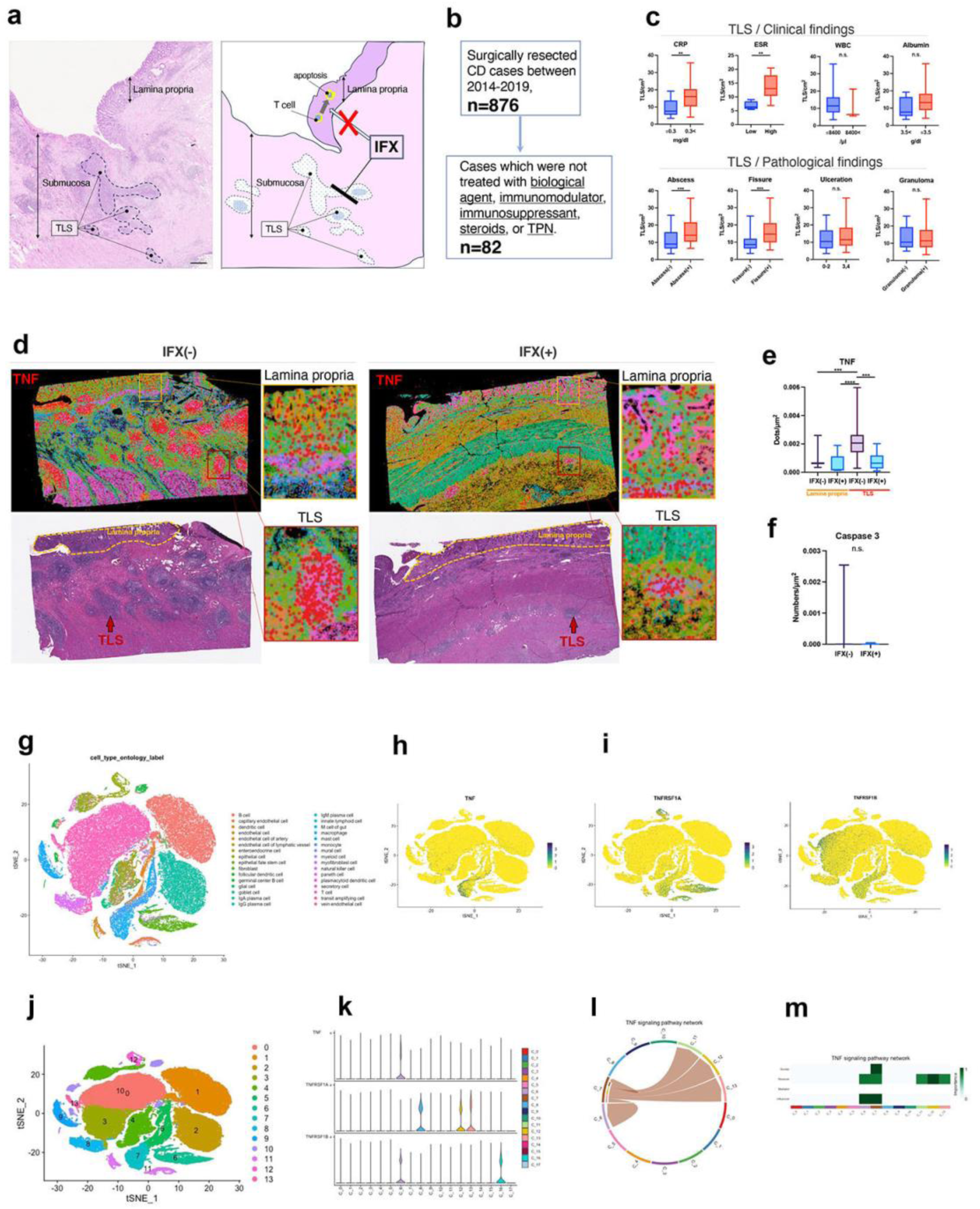
a Distribution of inflammation in Crohn’s disease tissue specimens (HE staining, schema). Theory of IFX mechanisms of action and hypotheses regarding the IFX mechanisms of action in the present study. Scale bar=500 μm. b Cases used to analyze the correlation between TLS and clinicopathological findings. c Association of clinical (CRP, ESR, WBC, Albumin) and histological (abscess, fissure, ulcer depth, and granuloma) findings with the number of TLS (n=82). d Distribution of TNF-expressing cells at mRNA level in the CD intestinal wall (IFX-, IFX+) by Xenium analysis. e Quantitative analysis of TNF levels in the TLS of IFX-treated and non-treated patients. f Quantitative analysis of the number of cleaved caspase 3-positive cells distributed in the TLS at the protein level using PhenoCycler analysis in IFX-treated- and non-treated patients. g Cluster analysis of single-cell RNA-seq data from untreated patients with Crohn’s disease (n=14). h Visualization of TNF-expressing cells by t-SNE projection. i Visualization of TNF receptors TNFR1 (TNFRSF1A) and TNFR2 (TNFRSF1B) using t-SNE projection. j Clustering analysis for CellChat analysis of TNF signaling. k Violin plot of clusters expressing TNF- and TNF-receptors. l Chord diagram for visualizing cell–cell communication through TNF signaling pathways, including TNF-TNFRSF1A and TNF-TNFRSF1B. m Heat maps of TNF signals contributing to outgoing or incoming TNF signaling in certain cell clusters.

The primary site of submucosal inflammation in patients with CD is the lymphoid follicle, often referred to as the tertiary lymphoid structure (TLS) (Fig. 1a). The TLS supports the local adaptive immune response and acts as an organized form of peripheral lymphoid tissue. It plays a crucial role in producing and regulating various cells involved in acquired immunity^6–8^. Several studies have highlighted the relationship between TLS and ulcerations, common lesions associated with CD. For instance, aphthous ulcers, the primary lesions of CD, are thought to develop directly above lymph follicles^9^. Additionally, our research indicates that plasma cells derived from TLS may play a role in intestinal ulceration by causing ischemia in the mucosa^10^. Furthermore, studies have shown that TLS can prevent ulcer formation in mouse models by inhibiting the development of colonic patches, a type of peripheral lymphoid tissue^11^. Therefore, it is possible that TLS, whose primary function is to mediate inflammation in CD, contributes directly to ulcer formation.

We hypothesized that if TLSs are the primary source of inflammation in CD and are directly linked to ulcer formation, then the primary target of inflammation-suppressing drugs, such as TNFα inhibitors, maybe the TLS rather than the immune cells infiltrating the lamina propria. Regarding TLS and the treatment of inflammatory diseases, some researchers have proposed that TLS could be a potential therapeutic target for patients resistant to conventional therapies^6^. In the context of inflammatory bowel disease, it is important to understand the evolution of TLS in response to treatment. However, there is still a lack of clarity regarding how TLS changes during treatment and whether it can be effectively targeted for therapeutic purposes^9^. TLS is mainly found in the deep intestinal tract in CD; therefore, changes in response to treatment have not been adequately analyzed in human specimens^9^. Our goal was to capture the changes in the TLS resulting from IFX administration. In this study, we conducted a detailed examination of alterations in TLS following IFX treatment, utilizing surgical cases that involved complete lesions in patients with CD. This study assessed the potential of the TLS as a therapeutic target.

## RESULT

### Correlation between TLS and clinicopathological findings of CD

TLS has been reported to be associated with disease activity in various inflammatory conditions^12^. However, to date, no study has clarified the link between TLS and CD. To investigate this association, we selected 82 of 876 surgical cases of patients with CD that had not undergone intensive therapies, such as biologics, steroids, or prolonged fasting. We only included cases that experienced minimal or no modifications from these therapies to compare TLS with the clinicopathological findings (Fig. 1b). TLS was more frequently observed in patients with high CRP levels and an increased erythrocyte sedimentation rate (ESR) (CRP, p=0.0026; ESR, p=0.0056; Fig. 1c). This was also noted more often in patients with low albumin levels, although the difference was not statistically significant. Regarding the pathological findings, TLS was more common in patients with abscesses and fissures (abscess, p=0.0008; fissure, p=0.0002; Fig. 1c), although there was no association with the presence of granulomas or depth of the ulcer. As described above, increased TLS formation was observed in CD with higher activity, a finding that supports the association between pathogenesis and TLS.

### Changes in TNF-expressing cells with or without IFX treatment and the cells involved in TNF signaling in CD

Using spatial analysis at the protein and mRNA levels, we analyzed several IFX-treated and treatment-naïve cases, and carried out a detailed analysis of the changes in TLS caused by IFX administration. Previous reports have identified macrophages and T cells (specifically Th1 cells and resident memory T cells) in the lamina propria with CD as sources of TNFα production^13,14^. However, there is a lack of studies confirming the distribution of TNFα-producing cells throughout the intestinal wall. To address this gap, we first evaluated the distribution of TNFα mRNA-expressing cells across the entire intestinal wall using surgical specimens that contained TLS. This assessment aimed to determine the specific locations within the intestinal wall where TNFα-producing cells, targeted by IFX, were located. TNF-expressing cells were primarily located within the TLS, and the expression of TNFα was significantly higher in the TLS compared to the lamina propria (p = 0.0003; Fig. 1d,e).

Additionally, in cases treated with IFX, the number of TNFα-expressing cells exhibited a decreasing trend, although there was no significant difference in expression within the lamina propria. In contrast, TNFα expression in the TLS was significantly reduced (p < 0.0001; Fig. 1d,e). Based on the mechanism of action of IFX, which induces apoptosis in TNFα-producing cells, it was hypothesized that these cells primarily undergo apoptosis in TLS rather than in the lamina propria. Therefore, we assessed the state of apoptosis in the TLS following IFX treatment using caspase 3 as a marker. No difference in the number of caspase 3-positive cells was observed between the IFX- and non-treated cases (Fig. 1f). This was thought to occur because most mTNFα-expressing cells were already undergoing apoptosis and had disappeared. IFX-treated patients had been receiving treatment for several months to years.

Next, single-cell analysis of cells expressing TNFα showed increased expression of TNF, mainly in macrophages (Fig. 1g,h). In contrast, TNFα receptors (TNFR1/TNFRSF1A, TNFR2/ TNFRSF1B) were expressed in various cells (Fig. 1g,i). CellChat analysis was conducted to identify the cells involved in TNF-signaling in patients with CD. Clusters 6 and 7, the latter being a cluster of macrophages, were identified as both receivers and influencers of this signaling. Additionally, cluster 7 acted as a sender. This indicates that macrophages transmit TNF signaling in both autocrine and paracrine manners and are associated with the pathogenesis of CD. Furthermore, the expression of BAFF, which is important for TLS formation, was upregulated in macrophages (Cluster 7), and Cellchat analysis showed that it interacted with B and plasma cells (Supplementary Fig. 1a,b,c,d,e,f). IFX also inhibited the interaction of TNF-producing cells with B and plasma cells, which are the main constituents of the TLS. Considering that IFX induced apoptosis of TNF-producing cells, we speculated that it inhibited the interaction between BAFF-positive macrophages and B cells and plasma cells.

### Changes in cells associated with TNF signaling under IFX treatment

Cells in Cluster 6, which are associated with TNF signaling, were analyzed, and we found that the expression of CXCL13, a cytokine important for the maintenance of lymphoid tissue formation, was increased (Fig. 2a,b). Identification of cells expressing CXCL13 in cluster 6 revealed that they were follicular dendritic cells (FDCs) and fibroblast reticulocytes (FRCs), which are distributed in the lymphoid tissue (Fig. 2c,d). The expression was higher in FDCs than in FRCs. Histological analysis showed a significant reduction in the expression of CXCL13 and its receptor, CXCR5, at both the protein and mRNA levels within the TLS in patients treated with IFX compared to those untreated (CXCL13^+^ cells, p<0.0001; CXCR5^+^CD20^+^ cells, p=0.004; Fig.2e,f; CXCL13 p<0.0001, CXCR5 p<0.0001; Fig. 2g,h). Analysis of cells expressing CXCL13 revealed no changes in FRC or Tfh in cases treated with IFX compared to those who were untreated cases; however, there was a significant decrease in FDC (CXCL13^+^CD21^+^ cells, p<0.0001; Fig.2i,j). A significant reduction in the expression of FDC marker molecules was also observed at the mRNA level (CD21, P <0.0001; Fig. 2k,l).

**Fig. 2.**
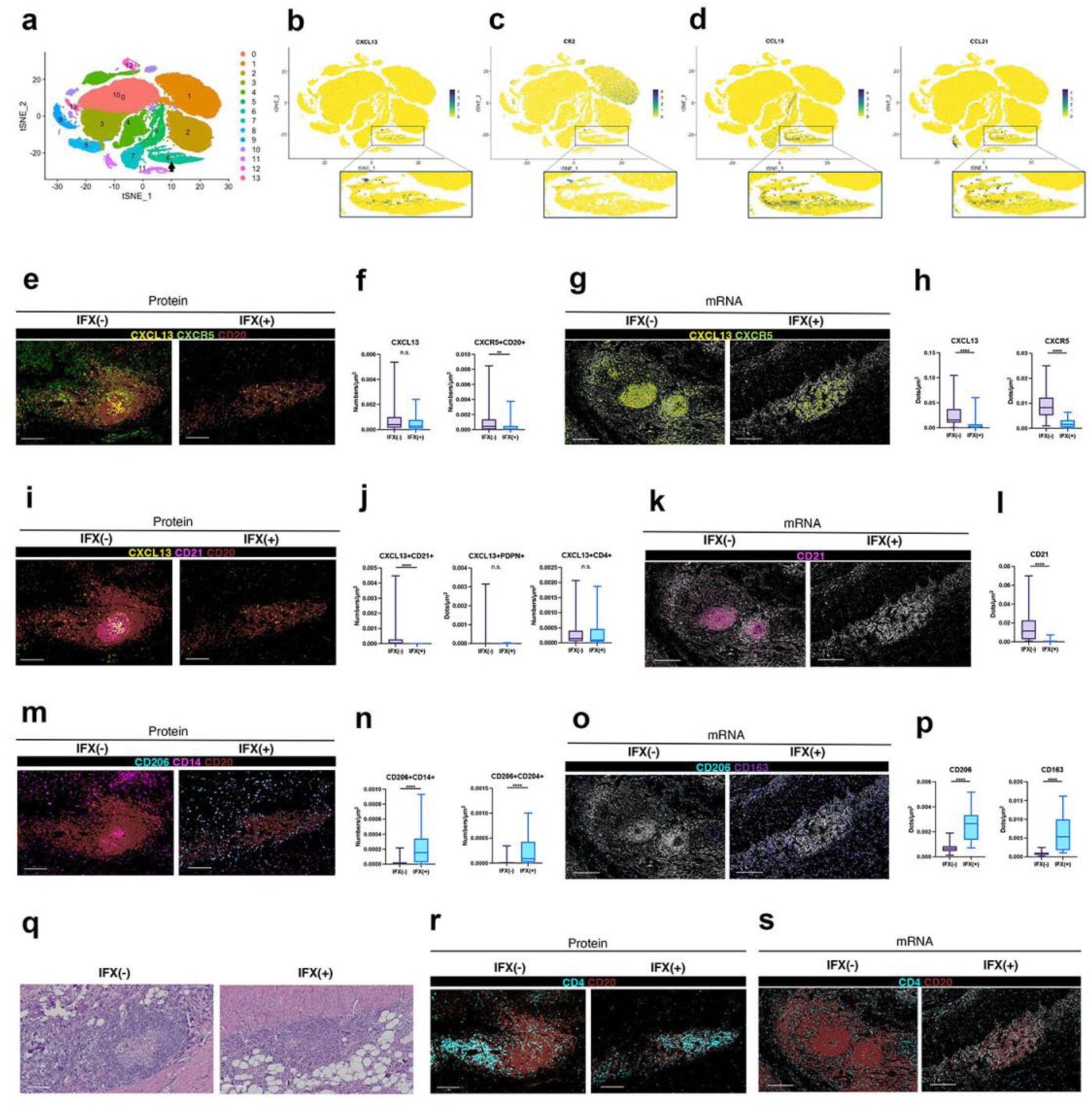
a Cluster analysis of single-cell RNA-seq data from untreated patients with Crohn’s disease (n=14). b Visualization of CXCL13-expressing cells by t-SNE projection. c Visualization of CR2-expressing cells by t-SNE projection. d Visualization of CCL19 and CCL21-expressing cells by t-SNE projection. e Multiplex immunohistochemistry of CXCL13, CXCR5 and CD20-positive cells in TLS in IFX- treated and non-treated cases. f Quantitative evaluation of CXCL13-expressing cells and CXCR5, CD20-positive cells at protein level in IFX-treated and non-treated cases. g Xenium spatial analysis of CXCL13, CXCR5 and CD20 positive cells within TLS in IFX treated and non-treated cases. h Quantified mRNA expression of CXCL13 and CXCR5 in TLS in IFX-treated and non-treated cases. i Multiplex immunohistochemistry of CXCL13, CD21 and CD20-positive cells in TLS in IFX-treated and non-treated cases. j Quantitative evaluation of CXCL13 and CD21 positive cells and CXCL13 and PDPN positive cells at protein level in IFX-treated and non-treated cases. k Xenium analysis of CD21 positive cells within TLS in IFX treated and non-treated cases. l Quantified mRNA expression of CD21 in the TLS in IFX-treated and non-treated cases. m Multiplex immunohistochemistry of CD206, CD14, and CD20-positive cells in TLS in IFX-treated and non-treated cases. n Quantitative evaluation of CD206 and CD14 positive cells, and CD206 and CD204 positive cells at protein level in IFX-treated and non-treated cases. o Xenium analysis of CD206 and CD163 positive cells within TLS in IFX treated and non-treated cases. p Quantified mRNA expression of CD206 and CD163 in TLS in IFX-treated and non-treated cases. q HE staining of TLS in IFX-treated and non-treated patients. r Multiplex immunohistochemistry of CD4 and CD20-positive cells in TLS in IFX-treated and non-treated cases. s, Xenium analysis of CD4 and CD20 positive cells within TLS in IFX treated and non-treated cases.

The TNF signaling pathway network analysis identified macrophages as Senders, Receivers, and Influencers. Moreover, mechanistically, the induction of M2 macrophages has been noted. Therefore, the characteristics of macrophages within the TLS were analyzed. CD204- and CD206-positive macrophages were increased (CD206+CD14+ cells, p<0.0001; CD206+CD204+ cells, p<0.0001; Fig. 2p,q), and the expression of molecules such as CD163 and CD206 were increased (CD163 p<0. 0001, CD206 p=<0.0001; Fig. 2r,s). In the analysis of 427 molecules conducted by Xenium, M2 macrophage-associated molecules, such as CD163 and MRC1, were among the most upregulated in patients treated with IFX. In contrast, molecules associated with lymphoid tissue formation, including CD21 and CXCL13, showed the most significant downregulation in these patients (Supplementary Fig. 2a,b).

Macrophages and T cells secrete various inflammatory cytokines, and IL6 and IL12 are therapeutic targets for inflammatory diseases^15,16^. Notably, IL6, IL12B, and IL1B were downregulated in IFX- treated patients (IL6: p<0.0001, IL12B: p<0.0001, IL1B: p<0.0001; Supplementary Fig. 3a,b), whereas the anti-inflammatory cytokine IL10 showed no difference in expression level between treatment with and without IFX (Supplementary Fig. 3c,d). These cytokine changes were thought to be associated with changes in the properties of M2 macrophages or apoptosis of TNF-expressing cells. Remodeling of the vascular system has also been reported as a function of TNF in lymphoid tissue formation^17^. When the vascular system was evaluated, vessels were distributed around the TLS and at the margins in the non-IFX-treated cases, whereas IFX treatment did not cause vessels to be distributed around the TLS but rather within the TLS structure (Fig. 2l). Quantitative assessment of blood vessels in the TLS showed increased expression of vascular molecules at both the protein and mRNA levels (CD31+ cells, p=0.0023; Supplementary Fig. 3e,f; CD34, p=0.0350; Supplementary Fig. 3g,h). Some vessels also express L-selectin ligand glycoproteins, collectively known as PNAd, which facilitate the migration of lymphocytes into lymphoid tissue. The number of high endothelial venules (HEV) expressing PNAd was significantly reduced in patients treated with IFX (CD31^+^MECA79^+^ cells p=0.0076 ; Supplementary Fig.3e,f).

Induction of regulatory T cell (Treg) via TNFR2 has also been reported as an effect of anti-TNFα antibodies, e.g. in rheumatoid arthritis^2^. Although the increase in Tregs is particularly controversial in human studies, in the present study, Tregs were more prevalent in IFX-treated TLS cases than in untreated cases (p=0.0218; Fig. 3i,j).

**Fig. 3.**
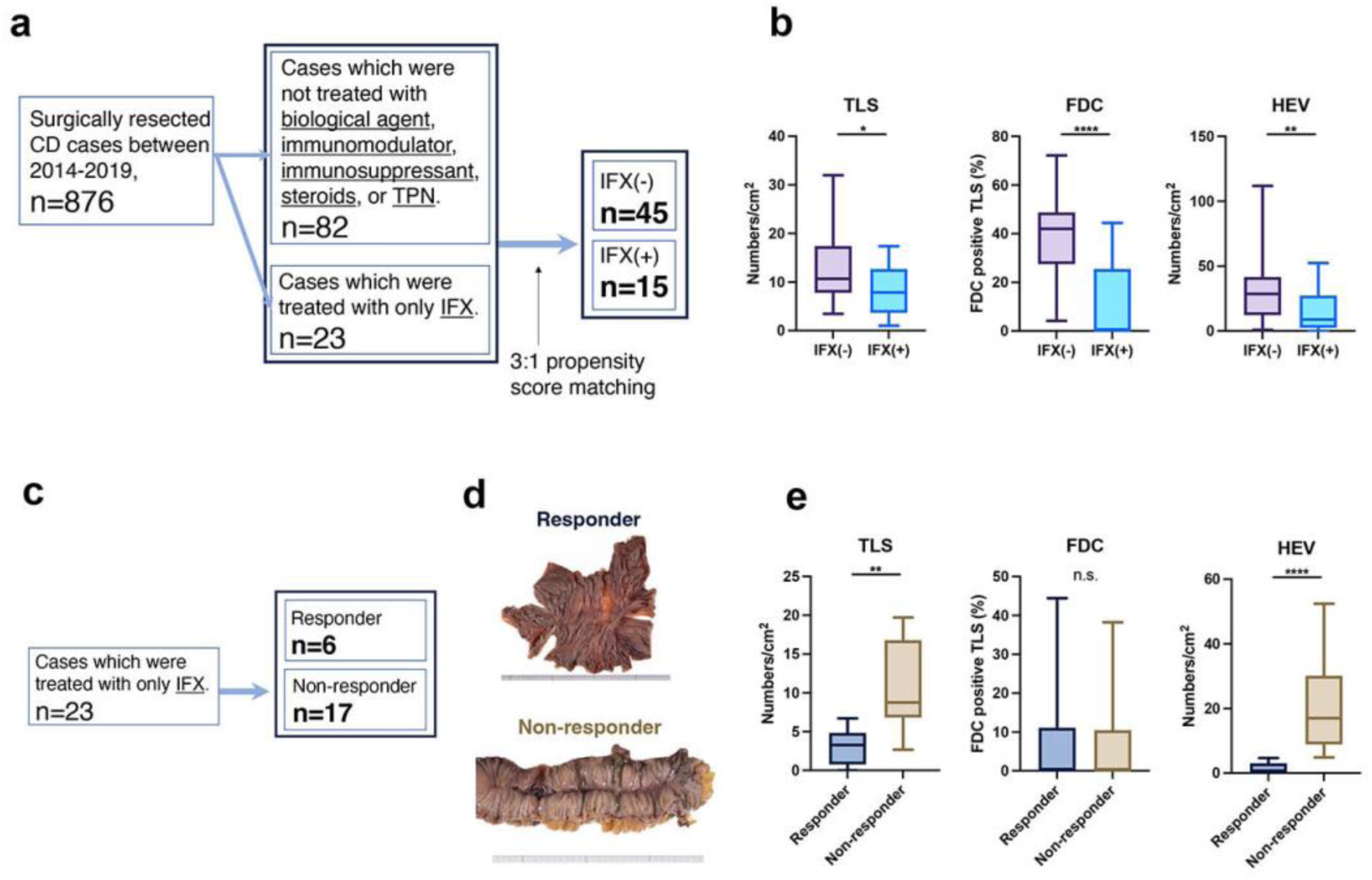
a Cases with covariate adjustment by propensity score matching (IFX-: n=45, IFX+: n=15). b Quantitative evaluation of TLS, FDC-positive TLS and HEV in covariate-adjusted IFX-treated and non-treated cases. c Responder and non-responder cases among those treated with IFX. d Macroscopic view of IFX responders and non-responders. e Quantitative assessment of TLS, FDC-positive TLS and HEV in IFX responder and non-responder

### Validation with a larger number of cases and differences between responder and non-responder

A study using 60 covariate-adjusted cases (45 not receiving IFX and 15 receiving IFX) employed a 3:1 propensity score-matching design to evaluate whether IFX treatment reduced TLS. In addition, the FDC and HEVs supplying lymphocytes to the TLS were assessed (Fig. 3a). In patients treated with IFX and TLS, the proportion of TLS with FDC and HEV was significantly reduced (TLS, p=0.0194; FDC, p<0.0001; HEV, p=0.0016; Fig. 3b). In addition, IFX-treated patients were classified as responders or non-responders, and their TLS status was assessed (Fig 3c). There were more TLS and HEV within the TLS in non-responders than in responders (TLS, p=0.0025; HEV, p=0.0001; Fig. 3d). However, the proportion of TLS with FDC was rarely observed in responders or non-responders, with no significant difference (Fig.3d). These results show that FDC in the TLS is almost completely lost in IFX-treated cases, whereas in non-responders, TLS and HEV remain, to some extent, even after the loss of FDC. The remaining HEV might not sufficiently disrupt the TLS; therefore, non-responders might not have experienced a reduction in inflammation.

### Expression of TLS signature genes before and after IFX treatment

Several molecules have been reported as TLS signature genes^18,19^. RNA-seq data from biopsy specimens before and after IFX administration were used to analyze changes in the expression of TLS signature genes. In responders, there was a significant reduction in the expression of TLS-related molecules before and after IFX treatment (NES: 1.765, nominal p-value = 0.0; Fig. 4a). In contrast, in the non-responders, no apparent changes in expression were observed before or after IFX treatment (Fig.4b). The expression of TLS signature genes was also significantly reduced before and after treatment with adalimumab, a TNFα inhibitor other than IFX, similar to IFX (NES: 1.512, nominal p value=0.004; Fig. 4c).

**Fig. 4.**
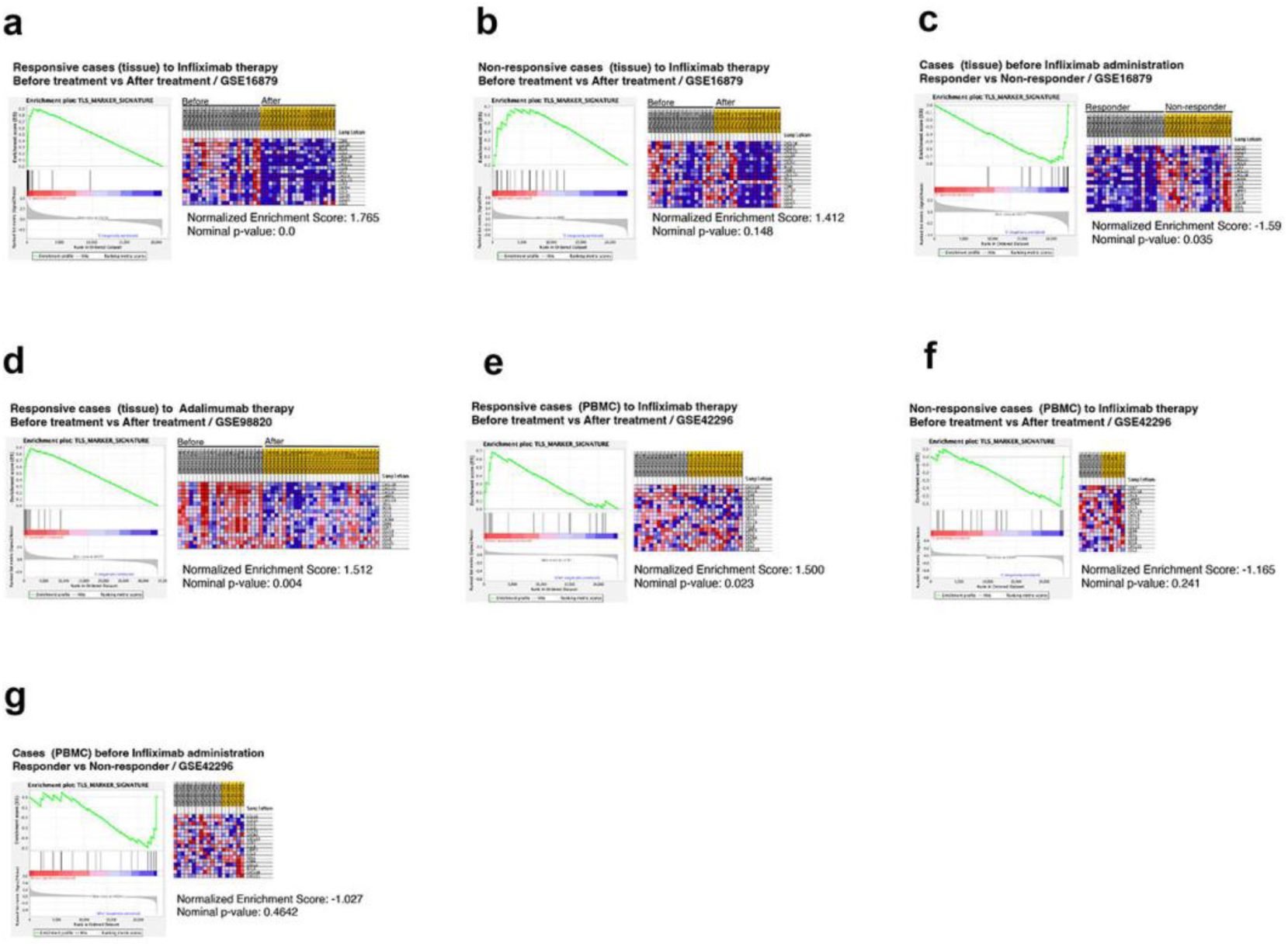
GSEA analysis of TLS signature gene set. a Enrichment plots in biopsy tissue from patients with CD with IFX response cases before and after IFX administration. b Enrichment plots in biopsy tissue from patients with CD with IFX non-responders before and after IFX administration. c Enrichment plots in biopsy tissue from patients with CD with IFX-responsive and IFX-resistant cases before IFX administration. d Enrichment plots in biopsy tissue from patients with CD with Adalimumab response cases before and after IFX administration. e Enrichment plots in PBMC from patients with CD with IFX response cases before and after IFX administration. f Enrichment plots in PBMC from patients with CD with IFX non-response cases before and after IFX administration. g Enrichment plots in PBMC from patients with CD with IFX-responsive and IFX-resistant cases before IFX administration.

A comparison of the expression of TLS signature genes before IFX administration between responders and non-responders revealed that responders had significantly lower expression levels of these genes (NES: 1.59, nominal p = 0.035; see Fig. 4d). This indicates that the TLS status could serve as a predictor of response to IFX treatment. Furthermore, the decreased expression of TLS-related molecules in responders was confirmed in both tissue samples and peripheral blood mononuclear cells (PBMCs) before and after IFX administration (Fig. 4e, f).

IFX treats several inflammatory diseases, including rheumatoid arthritis, ulcerative colitis, and CD^20^. We tested whether similar changes in the expression of TLS signature genes in ulcerative colitis and rheumatoid arthritis were observed in the samples before and after IFX treatment as in CD. In UC, the expression of TLS-related molecules decreased before and after IFX treatment in responders (NES: 1.646, nominal p-value =0.036; Fig.5a), whereas no apparent reduction was observed in non-responders (Fig.5b). Similar to CD, significant differences were found between responders and non-responders in the expression of TLS-related molecules before treatment (NES: 1.512, nominal p-value =0.007; Fig.5c). In rheumatoid arthritis, responder patients showed reduced expression of TLS signature genes both before and after treatment with etanercept, a type of TNFα inhibitor (NES: -1.79, nominal p value=0.004; Fig. 5d). In contrast, the non-responders did not show any reduction in gene expression (Fig. 5e). There was no difference in the expression of TLS-associated molecules between responders and non-responders before treatment with etanercept or certolizumab (Fig.5f), whereas the expression of TLS-associated molecules increased in non-responders treated with infliximab (NES: - 1.57, nominal p =0.029; Fig. 5g). These results suggest that TNFα inhibitors may target TLS not only in CD but also in non-CD inflammatory diseases such as ulcerative colitis and rheumatoid arthritis.

**Fig. 5.**
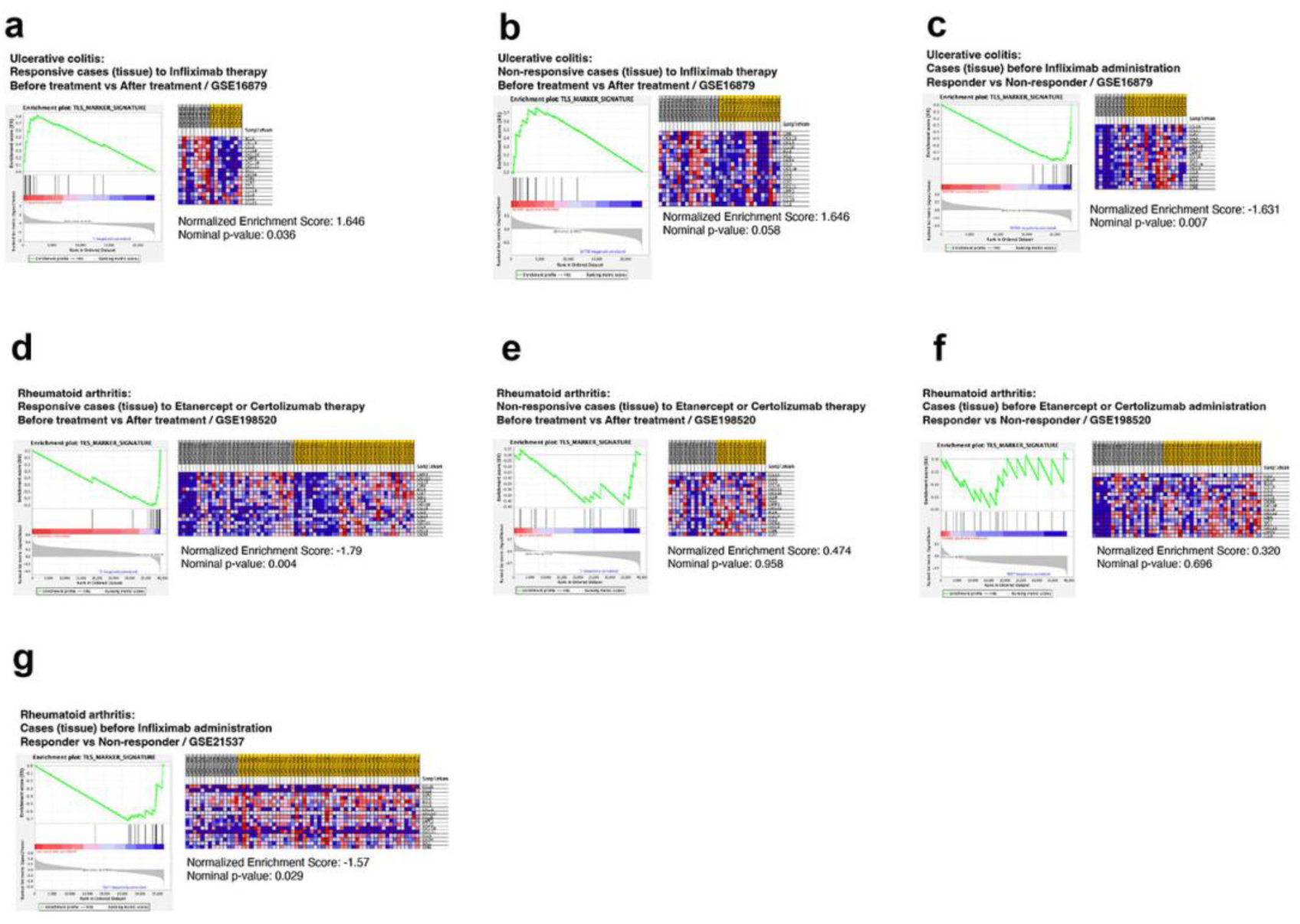
GSEA analysis of TLS signature gene set. a Enrichment plots in biopsy tissue from patients with ulcerative colitis with IFX response cases before and after IFX administration. b Enrichment plots in biopsy tissue from patients with ulcerative colitis with IFX non-response cases before and after IFX administration. c Enrichment plots in biopsy tissue from patients with ulcerative colitis with IFX- responsive and IFX-resistant cases before IFX administration. d Enrichment plots in biopsy tissue from patients with ulcerative colitis with Etanercept or Certolizumab response cases before and after Etanercept or Certolizumab administration. e Enrichment plots in biopsy tissue from patients with ulcerative colitis with Etanercept or Certolizumab non-response cases before and after Etanercept or Certolizumab administration. f Enrichment plots in biopsy tissue from patients with ulcerative colitis with Etanercept or Certolizumab-responsive and Etanercept or Certolizumab-resistant cases before Etanercept or Certolizumab administration. g Enrichment plots in biopsy tissue from patients with ulcerative colitis with IFX-responsive and IFX-resistant cases before IFX administration.

## DISCUSSION

The present study indicates that IFX, previously believed to act on T cells and macrophages in the lamina propria, disrupts the TLS by exerting various effects. Since TLS is linked to clinicopathological findings, and a direct causal relationship with ulcer formation has been proposed, IFX probably influences TLS to reduce inflammation. Furthermore, it was suggested that the state of the TLS could predict the therapeutic response to IFX in inflammatory bowel disease.

TNFα-producing cells, potential targets of IFX, were more abundantly distributed in the TLS than in the lamina propria and were significantly reduced with IFX treatment. Given the action of IFX, it was considered that IFX treatment caused TNFα-producing cells to undergo apoptosis. Although evaluation at the protein level was difficult because reliable antibodies for TNF were not available, it was thought that the concentration of TNF in the TLS was markedly reduced by IFX administration. Moreover, owing to the challenging issue of obtaining reliable antibodies against TNF, the effect of IFX on TNFα neutralization and apoptosis of TNF-producing cells was difficult to evaluate using protein expression analysis. TLS formation has been reported to be induced in TNFΔARE mice, which exhibit excessive TNF production, even without lymphoid tissue inducer cells (LTi cells)^21^. These mice present with impaired lymphatic flow associated with TLS formation, leading to spontaneous CD-like ileitis^22^. The formation of colonic patches, a type of peripheral lymphoid tissue, is promoted in a model of colitis induced by trinitrobenzene sulfonic acid. Treating these mice with LTβR-Ig, known to disrupt the formation of germinal centers like those found in the spleen, can also suppress the development of colonic patches and help prevent colitis. Considering that TLS is induced by TNF stimulation, that TLS can be associated with ulceration, that suppression of the colonic patch improves colitis, and that disruption of TLS is seen with TNF-targeted treatment, as shown in this study, this supports the theory that TLS is the main target of IFX.

Several effects of TNF on TLS formation have been reported^17,23^. The most important of these functions appears to be the induction and maintenance of FDC differentiation by TNF. TLSs form at various sites in the body owing to the differentiation of perivascular precursors, which are widely distributed, into FDCs when stimulated by TNF through TNFR1^23^. This study showed that CXCL13- positive FDCs almost disappeared in IFX-treated patients. However, this was thought to be due to the reduction and depletion of TNF in the TLS, which resulted in the disappearance of FDCs maintained by TNF stimulation. Such disruption of the FDC by TNF inhibitors has also been reported in rheumatoid arthritis^24^. Given that FDCs are the major secretors of CXCL13, which is essential for lymphoid tissue formation, it has been suggested that the changes that IFX exerts on FDCs are the main contributors to TLS disruption, and, therefore, are the main mechanism of action of IFX on CD.

TLS was more prevalent in non-responders to IFX than in responders, despite the almost complete disappearance of FDC. Therefore, it is possible that, in non-responders, there is a complementary mechanism that maintains TLS formation in the absence of FDC. In addition to FDC, Tfh and others produce CXCL13, and Tfh is also associated with TLS formation^25^. Considering that CXCL13+FDCs were markedly reduced under IFX treatment, whereas CXCL13+CD4 cells or CXCL13+FRC remained unchanged, this suggested that CXCL13-producing cells other than CXCL13+FDCs might have functioned complementarily in non-responders and maintained TLS formation. Indeed, IL12/23 inhibitors that block Tfh differentiation have been reported to be effective against IFX non-responders^26^. Therefore, the target of the IL12/23 inhibitors would be the TLS.

In IFX-naïve patients, the vessels were mainly distributed around the TLS and at the margins, whereas in IFX-treated patients, the vessels were distributed differently and embedded within the TLS. Vascular changes also included a decrease in HEV in IFX-treated patients. It has also been reported that TNF is associated with HEV induction^27^. Therefore, TNF inhibition likely reduces the number of HEV to some extent. However, when comparing responders and non-responders, significantly more HEV were found within the TLS of non-responders. Since TNF is associated with HEV induction, but LTBR-mediated stimulation, rather than TNF stimulation, has been reported to be important for the maturation and maintenance of HEV^28^, it is possible that in non-responders, a large number of already mature HEV are formed; consequently, TNF inhibition alone may not be sufficient to reduce HEV. Furthermore, the interpretation that residual HEV contributes to resistance to treatment with IFX is consistent with the reported success of α4β7 inhibitors in non-responders to IFX^29^. Given that HEV is a lymphoid tissue-specific vascular structure, it is suggested that TNFα and IL12/23 inhibition and α4β7 integrin inhibition targets the TLS.

The induction of M2 macrophages has been proposed as one of the direct mechanisms of action of IFX^2^, and M2 macrophages were significantly increased within the TLS in patients treated with IFX. It is not well understood how the increase in number of M2 macrophages affects TLS; however, there was also a decrease in pro-inflammatory cytokines with an increase in M2 macrophage number. Pro-inflammatory cytokines such as IL-6 act as regulators of TLS^30^. Therefore, increased levels of M2 macrophages and the associated changes in cytokine profiles may contribute to TLS disruption.

Predicting response to biological agents is a major challenge in treating inflammatory bowel disease^31^. In the present study, using an existing dataset of pre- and post-IFX biopsy specimens, the expression of TLS-related molecules before IFX administration was lower in responders than in non-responders. The Oncostatin M, IL-6 family cytokine has recently been reported useful for predicting CD activity and response to TNFα inhibitors^32^. Oncostatin M has also been implicated in TLS formation, and its increased expression promotes TLS formation^30,33^. Based on the results of this study and the discussion on Oncostatin M, TLS-related molecules might accurately predict the efficacy of TNFα inhibitory therapy. The TLS-associated molecules used in this study are a group of molecules that have been reported to be associated with cancer-related TLS, and further identification of TLS-associated molecules with responses to biologics specific to inflammatory diseases will be necessary. In addition, because most TLS-associated molecules are soluble, it is possible to assess IFX responsiveness by performing blood tests to investigate if the TLS status is related to treatment responsiveness.

Treatment with IFX for chronic inflammatory diseases such as ulcerative colitis and rheumatoid arthritis has also been shown to reduce the expression of molecules associated with TLS. Similar to CD, there was a correlation between TLS-related gene expression and treatment responsiveness under these conditions. IFX acts on various inflammatory diseases but is unlikely to act on each disease via entirely different mechanisms. TLS and ulceration have also been suggested as possible mechanisms in UC and CD. TLS is involved in the formation of lesions in CD and ulcerative colitis and rheumatoid arthritis, and IFX may have a therapeutic effect by disrupting TLS.

This study demonstrated that the primary target of IFX is the TLS by analyzing the changes in the entire intestinal wall resulting from IFX treatment. It also suggests that various biologics may target the TLS, revealing a new dimension in treating inflammatory diseases. Future investigations into the relationship between lesion formation and TLS in inflammatory diseases, along with treatment-induced changes in TLS, are anticipated to develop new treatments and methods for predicting treatment responses. This progress may pave the way for the development of precise personalized medicine for treating inflammatory diseases.

## MATERIAL AND METHOD

### Tissue selection and ethical approval

This study was approved by the Ethics Committee of the Yamate Medical Center (J-186). Medical records were reviewed to identify 876 cases of patients with Crohn’s disease who had undergone surgery between 2014 and 2019. From this group, 82 patients who did not receive intensive treatment, including biological agents, immunomodulators, or steroids, were selected to examine the association between TLS and clinicopathological findings. Propensity score matching was performed on these 82 cases and 23 cases treated with IFX alone. Analysis of tissue changes due to IFX was performed using 60 cases (45 without IFX and 15 with IFX) adjusted for covariates. Furthermore, 23 patients treated with IFX alone were classified into 6 responders and 17 non-responders, and histological differences between responders and non-responders were analyzed.

### In situ gene expression profiling (Xenium analysis)

Surgically resected specimens from patients with CD were fixed with 10% neutral buffered formalin and embedded in paraffin. Formalin-fixed paraffin embedded (FFPE) tissues were sliced to a thickness of 5 µm, trimmed to 8×4 mm, and specimens from 4 different samples (2 IFX treated samples / 2 treatment naïve samples) were mounted on xenium slide (10X Genomics) within sample areas (10.45 mm x 22.45 mm). The tissue sections were deparaffinized in xylene and rehydrated with ethanol. The cells were then treated with a permeabilization enzyme (CG000581, Rev A, 10x Genomics) for RNA accessibility. Hybridization probes were prepared using a pre-designed (Xenium Human Multi-Tissue Gene Expression panel) and custom (Xenium Custom Gene Expression panel, design ID: J89C6D) panels. Details of the probes are provided in Supplementary Table 1. The probes were hybridized overnight at 50 °C. A post-hybridization wash was conducted at 37 °C for 30 min, followed by ligation at 37 °C for 2 h and amplification at 30 °C for 2 h, according to the user guide. Autofluorescence quenching and nuclear staining were performed in the dark. Fluorescent probe hybridization and imaging were performed using a Xenium Analyzer (onboard analysis: version 1.1.0.2; software: version 1.1.2.4; 10x Genomics). Output images and expression profiles were analyzed using the Xenium Explorer (version 2, 10x Genomics). HE staining, along with MS4A1 and CD4 expression, was used as a reference to identify and annotate TLS. Samples were stained with H&E immediately after xenium decoding. Samples were washed in distilled water for 2 min and stained with hematoxylin for 15 min. Next, the sections were stained with a bluing reagent for 1 min, followed by eosin staining for 30 s.

### Multiplexed fluorescence immunostaining (PhenoCycler analysis)

Surgically resected specimens from patients with CD were fixed with 10% neutral buffered formalin and embedded in paraffin. FFPE tissues were sliced to a thickness of 6 μm, trimmed to 7×6 mm, and specimens from 8 different samples (4 IFX treated samples / 4 treatment naïve samples) were mounted on a hydrophilic slide glass (MAS-coat type, Matsunami glass Ind., Ltd.). Multiplexed fluorescence immunostaining was performed using the PhenoCycler system (Akoya Biosciences), following the manufacturer’s protocol outlined in the CODEX User Manual (Rev C). The samples were first dried using desiccant beads for 2 min, and then incubated in acetone for 10 min. Subsequently, they were placed in a humidity chamber for 2 min, followed by hydration using a Hydration Buffer. After this step, the samples were fixed using a Pre-Staining Fixing Solution to achieve a final concentration of 1.6% paraformaldehyde and were left to fix for 10 min. After washing the tissues with Hydration Buffer, the samples were incubated in Staining Buffer for 20 min. In a humidity chamber, the tissues were incubated with the antibody cocktail for 3 h. Details of the antibodies used are listed in Supplementary Table 2. After staining, the tissues were washed twice with the Staining Buffer and fixed with a Post-Staining Fixing Solution for 10 min. Subsequently, the tissues were washed thrice with 1 × PBS and incubated in ice-cold methanol for 5 min. They were then washed thrice with 1× PBS and fixed with a Final Fixative Solution for 20 min in a humidity chamber. The samples were washed three times with 1× PBS before being stored in Storage Buffer to prepare them for further assays. A Keyence microscopy system (BZ-X800, Keyence) controlled by a CODEX Instrument Manager (CIM, Akoya Biosciences) was utilized for multicycle fluorescence detection. The data were visualized and quantified using the HALO imaging software (Version 4.0). HE staining, along with CD20 and CD4 staining, was used as a reference to annotate TLS and quantitatively assess each cell within it.

### Propensity score matching

Characteristics differed between patients treated with IFX and those treated with minimal treatment. Propensity score matching analysis was performed to reduce bias between the groups. The propensity score was calculated using a logistic regression model that included the following variables: sex, Montreal classification, BMI, age, upper digestive tract lesions, anal lesions, CRP level, Harvey Bradshaw index, disease duration, smoking status, elemental diet, and 5ASA. A 1:3 nearest-neighbor matching technique was applied using the nearest-neighbor matching method with a caliper width of 0.3. All baseline characteristics were compared before and after propensity score matching. The Mann–Whitney U test was used for continuous variables, whereas categorical variables were analyzed using the chi-square test or Fisher’s exact test.

### Pathological evaluation of tertiary lymphoid structures in a propensity score matched larger cohort using immunohistochemistry

Surgically resected specimens from CD patients (15 patients treated with IFX and 45 patients not treated with IFX) were fixed with 10% neutral buffered formalin and embedded in paraffin. FFPE tissues were sliced to a thickness of 3 μm, and were mounted on a CREST slide glass (Matsunami Glass Ind., Ltd.). The slides were incubated at 70°C for a duration of 2 h, and slides were deparaffinized in xylene and rehydrated in graded ethanol solutions. Antigen retrieval was carried out at 97 C for 10 min with high pH (9.0) EDTA solution. Sections were incubated in the primary antibody overnight in 4 °C. The following primary antibodies were used: CD3 (1:400, CD3-12, Abcam, MA, USA). CD20 (1:200, L26, Agilent, CA, USA.), CD21 (1:500, 1F8, Abcam, MA, USA.), and MECA79 (1:50; MECA-79, Novus Biological, CO, USA). After washing the slides in phosphate-buffered saline with 0.005% Tween 20, they were incubated with a horseradish peroxidase-conjugated anti-rabbit or anti-mouse Ig polymer as a secondary antibody (Envision kit; Dako, Glostrup, Denmark) for 30 min at room temperature. The slides were treated with a streptavidin-peroxidase reagent and incubated in phosphate-buffered saline with 0.005% Tween 20. The samples were visualized by incubating the slides for 10 min in a solution containing 0.1 M Tris-HCl, 0.05% 3,3’-diaminobenzidine, 0.04% nickel chloride, and 0.0075% hydrogen peroxide (H_2_O_2_). The slides were counterstained with Mayer’s hematoxylin. All the slides were independently evaluated by two researchers (MK and YN). Aggregation of CD20-positive cells was assessed by TLS, with reference to hematoxylin and eosin (HE) staining and CD3 staining. TLS with CD21 positivity within the TLS was defined as FDC- positive TLS and CD21-negative. For HEV, one MECA79-positive location was considered one HEV.

### Single cell RNA-seq

scRNA-seq data were obtained from the publicly available Single Cell Portal (https://singlecell.broadinstitute.org/single_cell/study/SCP1423/predict-2021-paper-cd)^34^. This dataset includes 14 untreated pediatric Crohn’s disease samples, analyzed for 22431 gene expressions across 107432 cells. The gene expression matrix was imported into the R software version 4.3.3 and converted into Seurat objects using the Seurat package (version 5.1.0). First, gene expression count data were normalized using the NormalizeData function and scaled using the ScaleData function. Next, the FindVariableFeatures function was employed to select the top 2,000 variable genes, followed by principal component analysis (PCA). Cell clusters were identified using the FindNeighbors (dimensions = 10) and FindClusters functions (resolution = 0.3). To visualize the data in two dimensions, t-SNE (t-distributed Stochastic Neighbor Embedding) was performed. Cell-cell interaction analysis was conducted for Clusters 6 and 7 (macrophages) in relation to other cell types using the R package CellChat version 1.6.1, which utilizes a public built-in database for ligand-receptor interactions^35^.

### Microarray analysis

Microarray data for CD and UC before and after IFX administration (GSE16879) were downloaded from GEO. This dataset comprised biopsy specimens taken from the active sites of 24 cases of ulcerative colitis and 37 cases of CD before and 4-6 weeks after the first IFX administration. The response to IFX was assessed based on endoscopic and histological findings at 4-6 weeks after IFX administration. A dataset using adalimumab for patients with CD was downloaded (GSE98820). This dataset contained 76 biopsies from 10 patients. All the patients were responders. PBMC data from patients with CD were downloaded (GSE42296). This data included 20 CD specimens; 14 were responders and 6 were non-responders. A dataset of rheumatoid arthritis synovium was downloaded (GSE198520) containing synovial samples from 46 patients with rheumatoid arthritis treated with TNFα inhibitor therapy (19 cases: Etanercept, 27 cases: Certolizumab). These tissues were collected before and 12 weeks after TNFα inhibitor treatment. The dataset of patients with RA treated with IFX was downloaded (GSE21537). These data sets included the synovial membranes of 62 patients: 18 responders, 30 moderately responders, and 14 non-responders. Responsive and moderately responsive cases were analyzed as responders.

### Gene Set Enrichment Analysis

Gene Set Enrichment Analysis (GSEA) was performed using GSEA software (v.4.3.2) from the Broad Institute (Massachusetts Institute of Technology). The differential expression of 17 molecules known as TLS signature or marker molecules was evaluated among responders and non-responders across various diseases treated with TNFα inhibitors. The molecules assessed included CXCL9, CXCL10, CXCL11, CXCR4, CXCL13, CCL18, CCL19, CCL21, BCL6, CCR7, CD86, CCL3, SELL, LAMP3, CCL8, CCL5, and CCL2^18,19^. GSEA evaluates genes in a ranked list from highest to lowest to calculate the enrichment score (ES) for a specific pathway. ES increases when a gene is part of a pathway and decreases when it is not. The normalized enrichment score (NES), which indicates how enriched a pathway is within the ranked list, was determined by the maximum value of the running sum normalized according to pathway size. A positive NES indicates enrichment at the top of the list, whereas a negative NES indicates enrichment at the bottom. A nominal p-value of ≤ 0.05 was considered statistically significant in the enrichment plots.

## Acknowledgements

This study was supported by JSPS KAKENHI Grant Number JP22H04925 (PAGS). We thank Takeshi Watanabe for carefully reading and giving critical comments on this manuscript.

## Contributions

M.K. conceived and designed the study. Y.N., S.O., A.F., T.K., T.Y., and M.K. generated and collected data. Y. S., S. O., S. N., and M. K. analyzed the data. Y.N., S.N., Y.S., and M.K. wrote the manuscript. Y. N. and M. K. prepared the figures. Y.N., K.Y., K.Y., K.O., S.N., Y.S., T.Y., K.A., T.O. and M.K. edited the manuscript. All authors have read the manuscript and agree with its contents.

---

**Supplementary Fig. 1.**
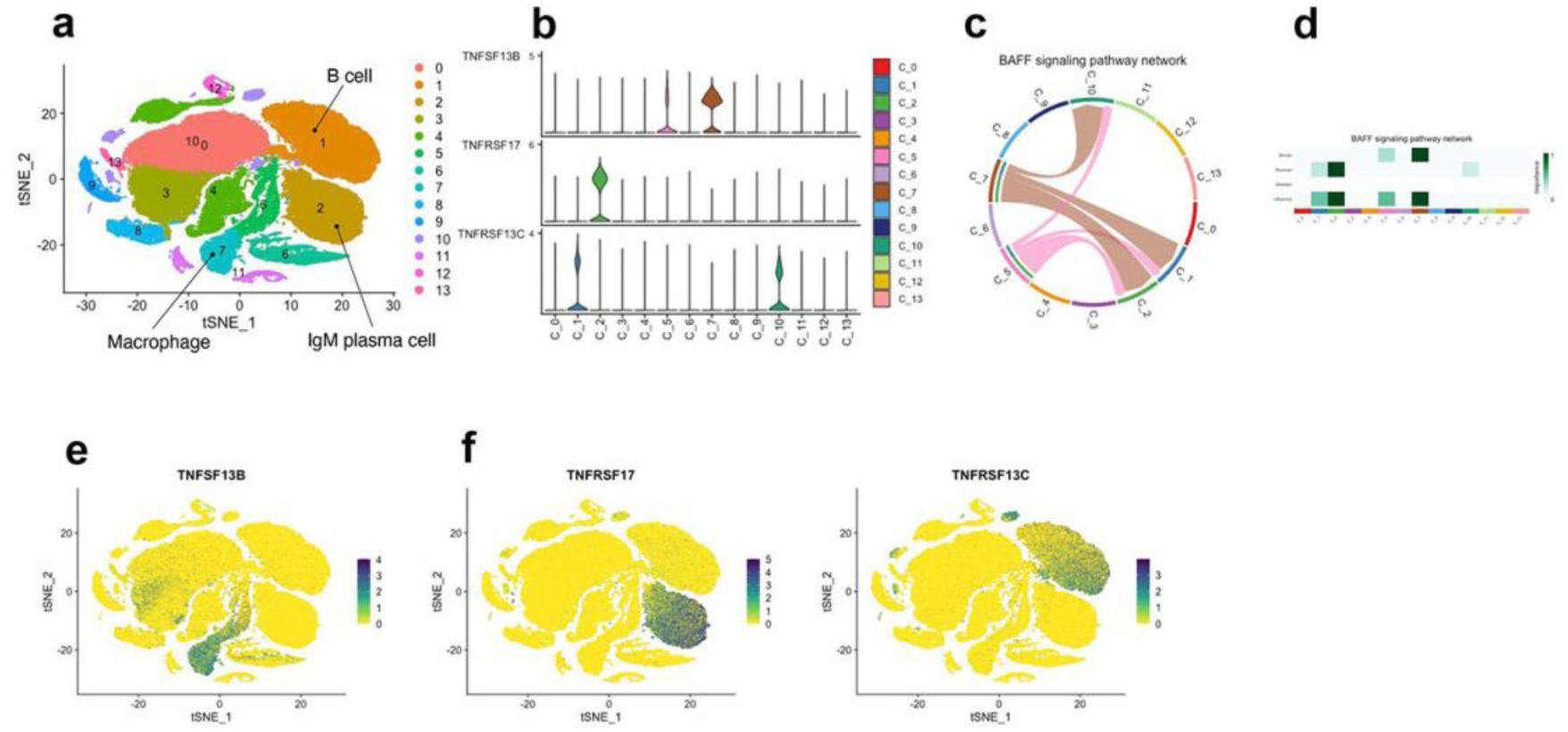
CellChat analysis of BAFF signaling pathway in CD. a Clustering analysis of single-cell RNA-seq data from untreated patients with Crohn’s disease (n=14). b Violin plot of clusters expressing TNFSF13B, TNFSF17 and TNFRSD13C. c Chord diagram for visualizing cell-cell communication through BAFF signaling pathways, including TNFSF13B, TNFSF17 and TNFRSD13C. d Heatmaps of TNF signals contributing to outgoing or incoming BAFF signaling of certain cell clusters. e Visualization of the TNFFSF13B by t-SNE projection. f Visualization of the TNFRSF17 and TNFRSD13C by t-SNE projection.

**Supplementary Fig. 2.**
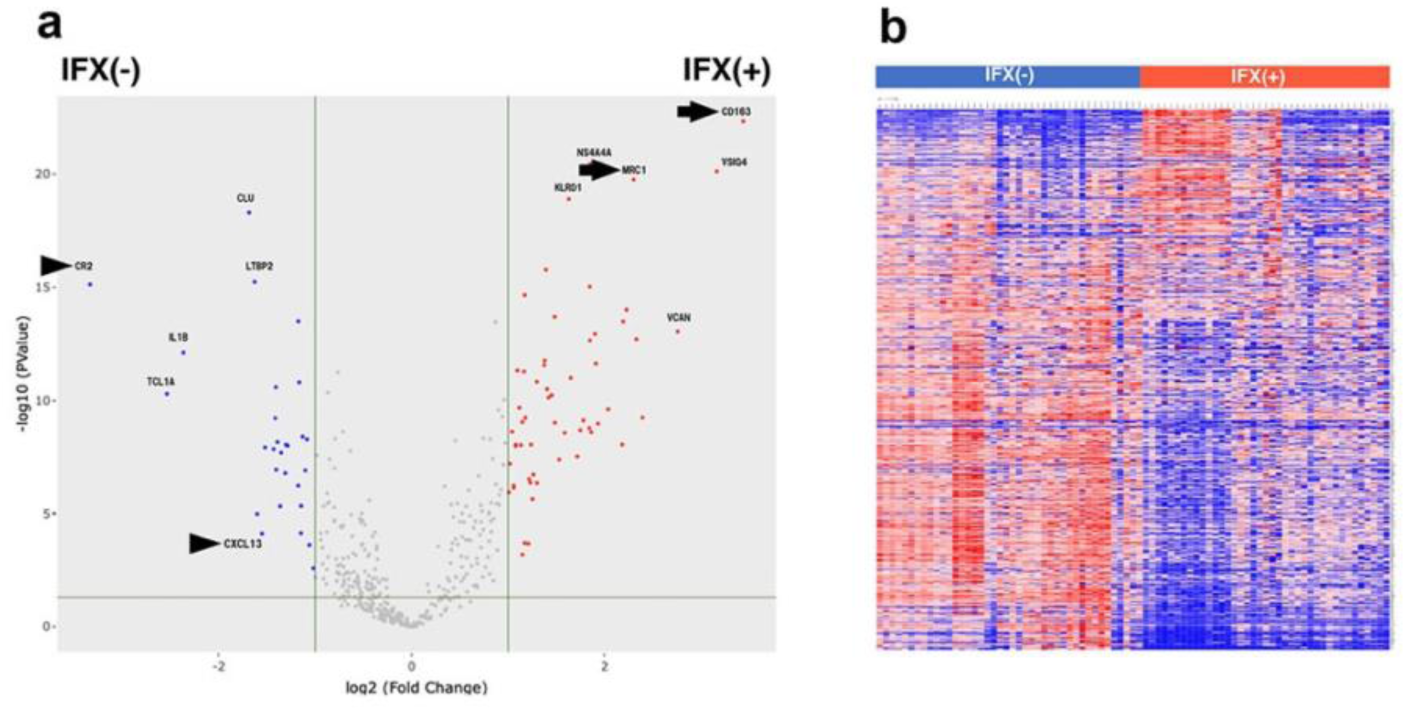
Comparison of expressions of 427 molecules in the TLS of IFX-treated and non-treated patients by Xenium analysis. a,b Heatmap and Volcano plot showing global changes of IFX administration. For Volcano plots, a p value < 0.05 and a − 1.5 > fold change > 1.5 were used as threshold.

**Supplementary Fig. 3.**
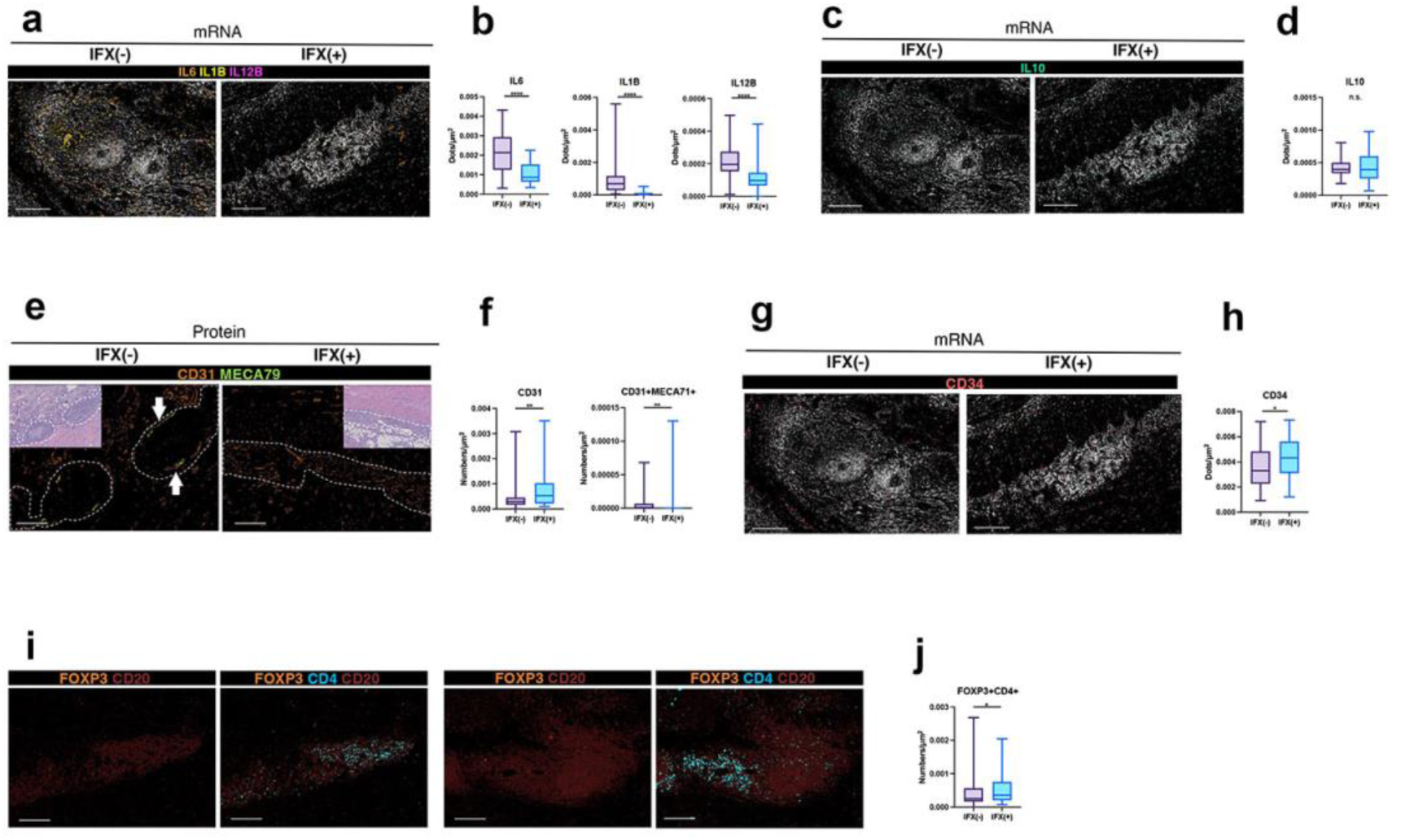
A Xenium spatial analysis of IL6, IL1B, and IL12B positive cells within TLS in IFX treated and non- treated cases. b Quantified mRNA expression of IL6, IL1B, and IL12B in TLS in IFX-treated and non- treated cases. C Xenium spatial analysis of IL10 positive cells within TLS in IFX treated and non- treated cases. d Quantified mRNA expression of IL10 in TLS in IFX-treated and non-treated cases. e Multiplex immunohistochemistry of CD31 and MECA79-positive cells in TLS in IFX-treated and non-treated cases. f Quantitative evaluation of CD31 and MECA79 at protein level in IFX-treated and non-treated cases. g Xenium spatial analysis of CD34 positive cells within TLS in IFX treated and non-treated cases. h Quantified mRNA expression of CD34 in TLS in IFX-treated and non-treated cases. i Multiplex immunohistochemistry of FOXP3, CD4, and CD20-positive cells in TLS in IFX- treated and non-treated cases. j Quantitative evaluation of FOXP3 and CD4 at protein level in IFX- treated and non-treated cases.

**Supplementary Fig. 4.**
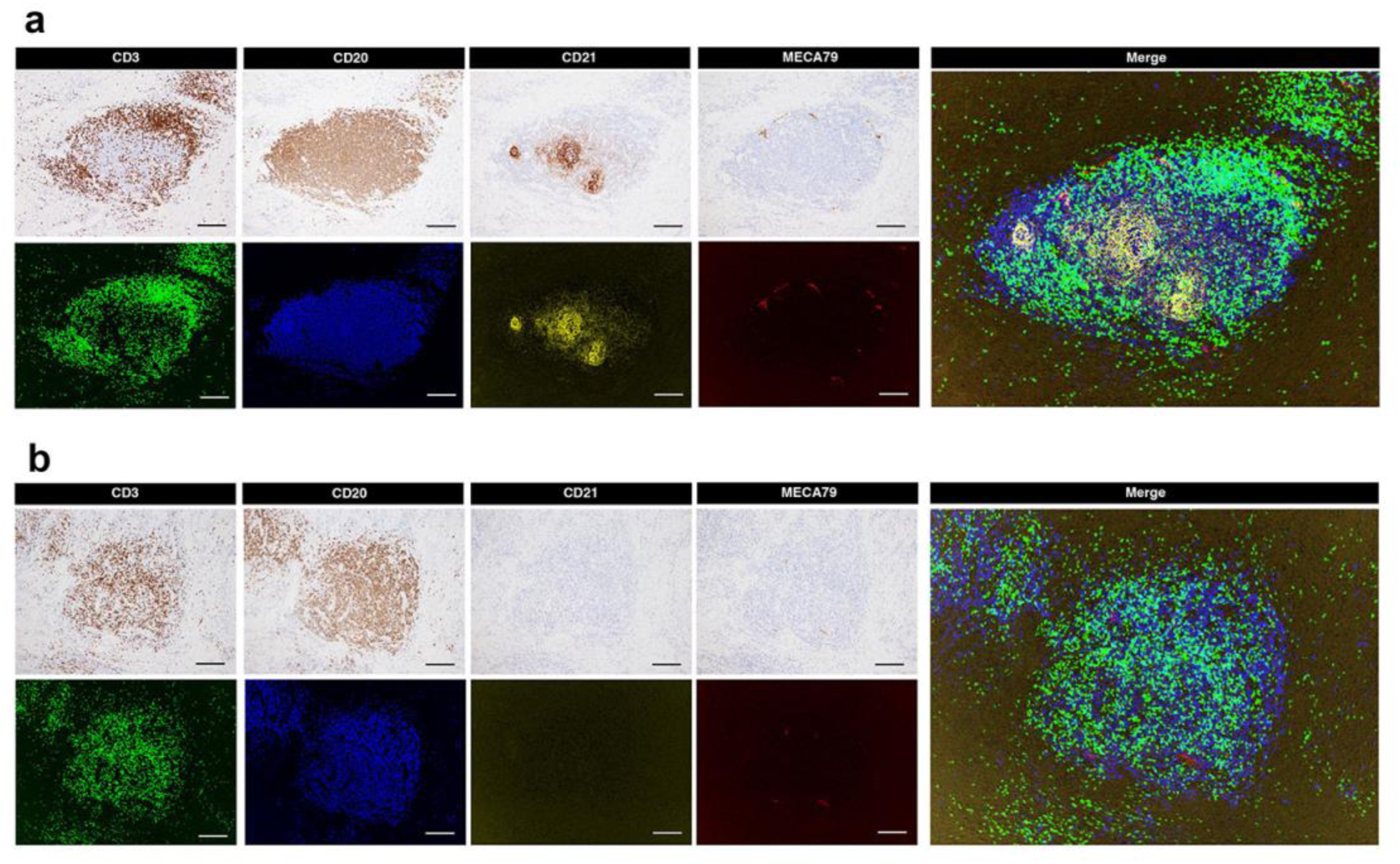
a Representative image of immunostaining for CD3, CD20, CD21, and MECA79 in treatment naïve cases. b Representative image of immunostaining for CD3, CD20, CD21, and MECA79 in cases treated with infliximab.

**Supplementary Fig. 5.**
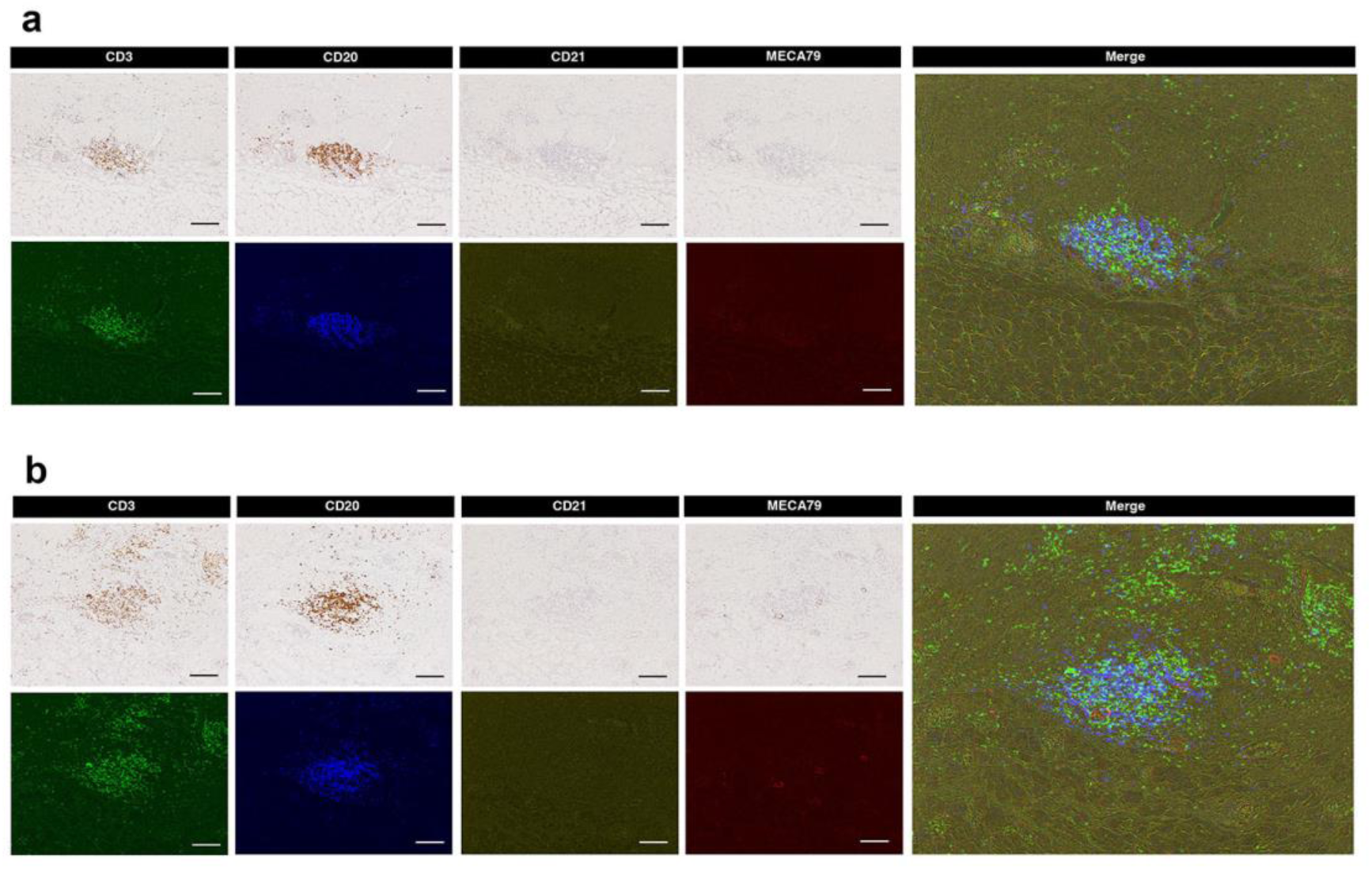
a Representative image of immunostaining for CD3, CD20, CD21, and MECA79 in IFX responsive cases. b Representative image of immunostaining for CD3, CD20, CD21, and MECA79 in IFX non- responsive cases.

